# Exploratory positional differences in observed CDRH2 lineage-state transitions following HIV-1 germline-targeting vaccination

**DOI:** 10.64898/2026.11.07.765060

**Authors:** Kazi Tanzina Begum, Kazi Abdul Gofur Rafea

**Author notes:** Corresponding author *Email address:* (Kazi Tanzina Begum).

## Abstract

**Background:** Germline-targeting HIV-1 vaccination recruits VRC01-class B-cell precursors, but the positional patterns of observed longitudinal B-cell receptor lineage-state changes remain incompletely described. We compared CDRH2 profiles between homologous and heterologous boosting regimens using published IAVI G002 data.

**Methods:** Participant-specific Week-8 lineages were prospectively matched to Week-24 observations. Germline-to-nongermline changes in sampled lineage-modal residues were evaluated at seven IGHV1-2*02 CDRH2 positions using participant-balanced, mean-centered profiles and exact participant-label permutation inference. Sampling-depth and sorting-probe sensitivity analyses assessed observability. A separate participant-held-out benchmark evaluated antibody-language-model representations for predicting observed Week-24 lineage detection.

**Results:** The CDRH2 analysis included 332 lineages, 15 participants, 2,179 eligible lineage-position observations, and 241 observed state changes. The regional profile statistic was 0.027066 (exact p = 0.001709; six-region adjusted q = 0.010256), with descriptive differences at V-relative positions 51, 54, and 55. Most eligible longitudinal observations had one assessable cell per visit. In seven Group-1 participants, Week-24 dual-probe sampling recovered 181 baseline lineages with eOD-GT8 and 143 with Cg28v2, including 60 detected by both probes. In a separate cohort of 2,739 lineages from 17 participants, IgBERT and AbLang-2 representations did not improve participant-balanced average precision over a four-feature biological baseline.

**Conclusions:** Observed CDRH2 lineage-modal-state profiles differed between regimens in this exploratory secondary analysis. Sparse sampling, potentially probe-related ascertainment, and absent linked functional validation preclude inference about mutation timing, functional improvement, or causal booster effects.

## 1. Introduction

Original G002 study relationship. Willis et al. [1] described HIV-1 germline-targeting vaccine-induced VRC01-class antibody maturation, including key heavy chain somatic mutations and functional antibody profiles. We do not claim any new such biological findings. Our unique analytical question is if observed Week 8 to Week 24 lineage modal amino acid state acquisitions, across 7 selected IGHV1-2*02 CDRH2 positions, differ between homologous and heterologous boosting groups, using participant-balanced inference. We did not preregister a set of confirmatory positions. Our selected 7 positions were based on the recorded data support criteria. We address the spatial pattern of observed state acquisitions, not increased total mutation, increased binding, or better vaccine efficacy. HIV-1 germline-targeting vaccines aim to recruit precursors of broadly neutralizing antibodies (bnAbs) and then guide their maturation through sequential immunizations. Human priming studies have shown VRC01-class precursor recruitment [2]. Preclinical and germinal center studies provide rationale for heterologous boosting and the role of precursor frequency and antigen affinity [3,4]. Sampling depth, antigen sorting probes, and complementary functional antibody measurements are all important for interpreting vaccine-induced BCR repertoires accurately [10]. The randomized IAVI G002 trial tested mRNA–lipid nanoparticle immunogens expressing eOD-GT8 60mer and core-g28v2 60mer [1]. Here, we study two groups: Group 1 received eOD-GT8 at Weeks 0 and 8 (homologous boost); Group 2 received eOD-GT8 at Week 0 and core-g28v2 at Week 8 (heterologous boost). Our longitudinal BCR comparison utilizes Week 8 as the post-prime reference visit (not unvaccinated baseline) and Week 24 as the post-boost follow-up.

Regimen-related VRC01-class maturation differences, including participantlevel 90th percentile counts of key heavy chain and HCDR2 residues, were previously established by Willis et al. (2025) (Figure 5E–F; Table S61). Position-specific heavy chain key residue patterns were shown across groups and timepoints (Figure S39) [1]. We do not claim any new such general findings. Our secondary analysis, instead, explores the participant-centered positional shape of observed Week 8 to Week 24 germline to non-germline lineage modal state changes, in prospectively defined and subsequently recovered IGHV1-2*02 lineages. This is a unique exploratory endpoint that complements the original analysis without demonstrating mutation timing, functional improvement, or a causal booster effect. We do not assume V-relative positions are equivalent to standardized antibody residue numbering. We compare participant-weighted, within-region-centered positional profiles (as opposed to cells or lineage-position observations as independent trial participants). We also assess longitudinal cell depth support and a same-visit dual probe sampling contrast within Group 1. These help characterize what is observable in the repertoires; they do not separate vaccination biology from antigen-dependent ascertainment or determine mutation timing. Finally, a separate participant-held-out benchmark asks if pretrained antibody sequence representations add predictive value (above simple Week 8 biological features) for *observed* Week 24 lineage recovery. This secondary prediction label is not the CDRH2 mutation state endpoint [7–9].

## 2. Results

### 2.1. Week 8 lineage reconstruction verification

The frozen G002 coverage audit included 32 Group 1 and Group 2 participants, and 5,579 qualifying post-prime Week 8 eOD-GT8-sorted B cells. Eighteen participants (8 Group 1; 10 Group 2) provided 3,894 eligible IGHV1-2*02 Week 8 baseline lineages. Seventeen had qualifying Week 24 IGHG cells, Cg28v2-sorted, VRC01-class, for 3,868 baseline target lineages. One Group 1 participant (G002-479, 26 target lineages) had no qualifying Week 24 source cells. Two Group 2 participants (G002-947 and G002-957) had source cells, but no eligible matched target lineages (nondetection does not indicate lineage loss). 15 participants (7 Group 1; 8 Group 2) had 337 detected Week 24 target lineages, of which 332 entered the mutation state analysis (Figure 2). Thus, our CDRH2 comparison is on selected participants with evaluable longitudinal lineage observations, not all vaccinated participants.

| Analytical cohort stage | Group 1 | Group 2 | Total |
|---|---|---|---|
| Participants in coverage audit | 15 | 17 | 32 |
| Participants with eligible IGHV1-2*02 target lineages at Week 8 | 8 | 10 | 18 |
| Participants with qualifying Week 24 source cells | 7 | 10 | 17 |
| Participants with matched Week 24 lineages and assessable CDRH2 profiles | 7 | 8 | 15 |

We compared reconstructed Week 8 clusters with the provided author-cluster reference for 5,579 shared cells to assess computational lineage definition concordance. The partitions agree for 5,567 cells (2,585/2,593 Group 1; 2,982/2,986 Group 2). 12 cells are discordant. Adjusted Rand index is 0.994938. This indicates high concordance with an existing annotation on shared cells, not proof of biological clonal ground truth. An exploratory group comparison of participant-level lineage detection fractions shows a Group 2 minus Group 1 mean difference of 0.002549 (exact two-sided p = 0.932024). This does not indicate equivalence or rule out sorting or ascertainment-related selection bias. The original target lineage coverage audit and the separate model prediction cohort use different eligibility criteria and denominators. Thus, we report these populations separately below.

### 2.2. Participant-level CDRH2 positional state profile differences

Our original 7-position analysis shows differences in observed CDRH2 lineage modal state profiles between groups for 15 participants (7 Group 1; 8 Group 2). The centered profile statistic is 0.027066, with exact p = 0.001709 and BH-adjusted q = 0.010256 (across 6 regional comparisons). Participant omission p-values range from 0.001166 to 0.003996. Position omission p-values range from 0.001399 to 0.003108. We then performed a coordinate-defined 8-position sensitivity analysis using the same participants. The statistic is 0.023722, with exact p = 0.001709. Participant omission p-values range from 0.001166 to 0.003996. These analyses reuse the study cohort and do not provide independent validation. The original 6-region adjusted q-value is not an adjusted q-value for the subsequent 8-position analysis. Our corresponding Week 8 post-prime profile comparison shows p = 0.631702. A non-significant Week 8 post-prime comparison does not indicate equivalence.

The profile difference is characterized by higher acquisition fractions at V-relative position 51 in Group 1, and positions 54 and 55 in Group 2. Equally weighted participant means are 47.5% vs 16.1% (position 51), 10.6% vs 29.1% (position 54), 0.0% vs 23.4% (position 55) (Group 1 vs Group 2). No acquisition is observed at position 55 among 138 eligible Group 1 lineage-position observations, or among 7 Group 1 participants. Group 2 shows 45 acquisitions among 182 eligible observations, across 6 of 8 participants. These are descriptive position-specific findings, not separately tested hypotheses. The largest centered group contrasts are negative (position 51) and positive (positions 54 and 55). These directions are maintained in the 8-position analysis. Position-specific summaries are descriptive. The reported permutation p-value is on the overall centered profile, not separate hypotheses at individual positions. Mutation fractions are conditional on detected lineages and position-specific assessability.

Figure 1 shows participant-level CDRH2 positional acquisition profiles and the 6-region comparison.

**Figure 1:**
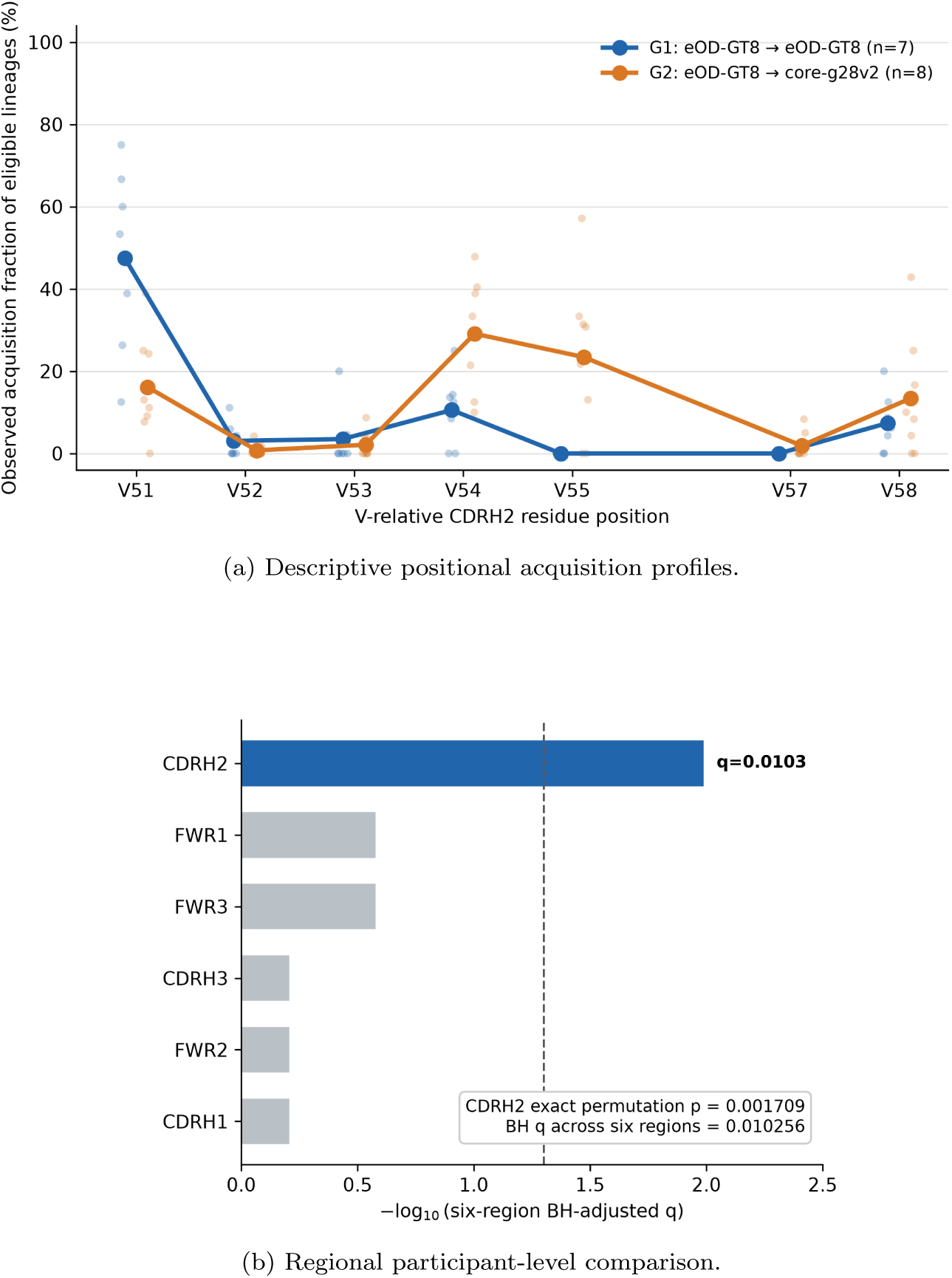
Exploratory observed G002 CDRH2 lineage modal state transitions. (A) Equally participant-weighted fractions at 7 selected IGHV1-2*02 V-relative positions (51, 52, 53, 54, 55, 57, 58), post-prime Week 8 to post-boost Week 24. (Group 1, *n* = 7; Group 2, *n* = 8). Faint points indicate participants. (B) Across region comparison of participant-centered positional profiles. CDRH2 exact participant-label permutation *p* = 0.001709 and 6-region BH-adjusted *q* = 0.010256 apply to the overall regional profile, not individual positions. Sampling-qualified observed states do not indicate mutation timing, fixation, causal booster effects, or function.

**Figure 2:**
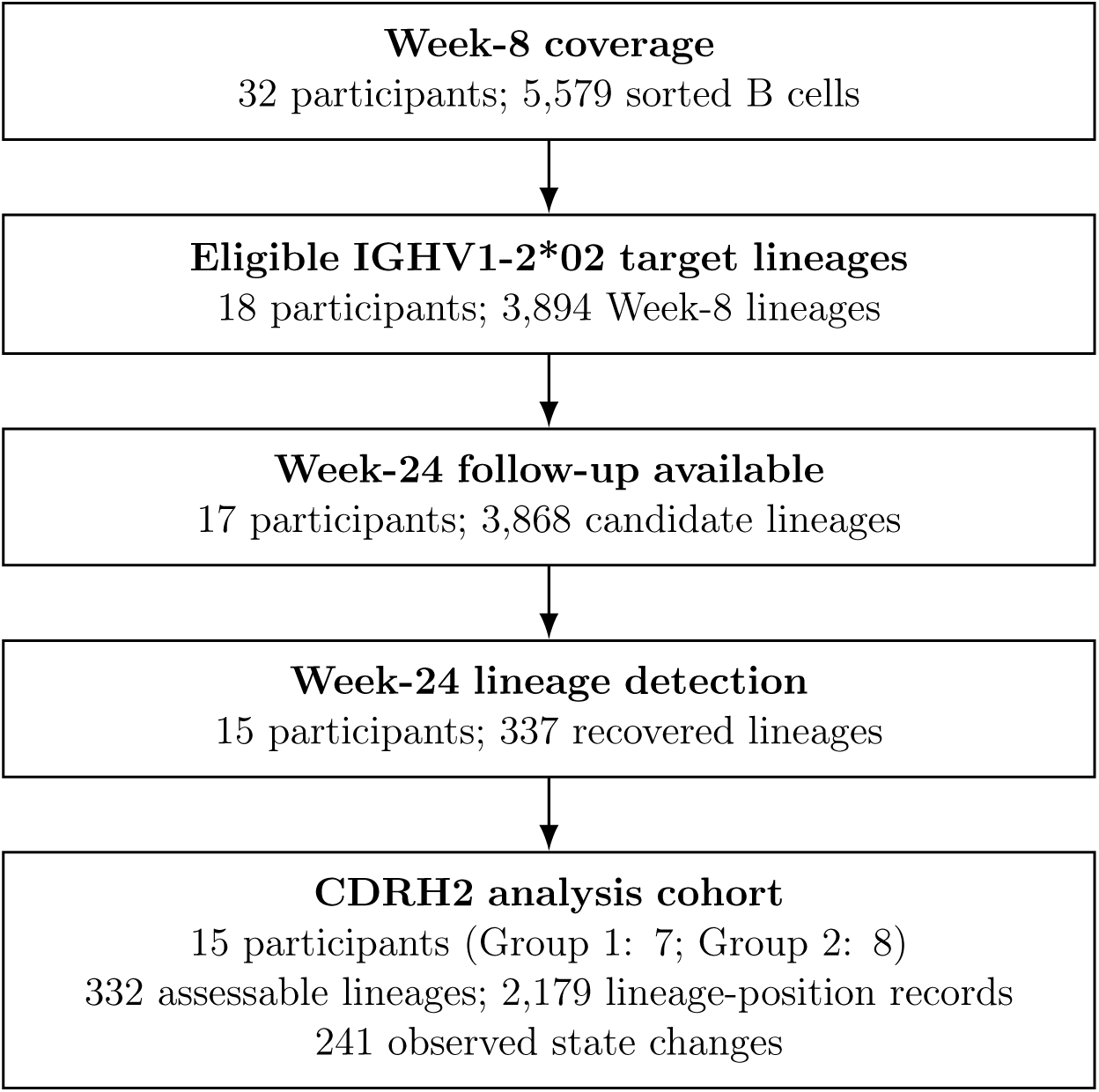
G002 longitudinal analytical cohort and eligibility flow. From 32 Group 1/Group 2 participants and 5,579 Week 8 cells, 18 participants provide 3,894 eligible IGHV1-2*02 target lineages. 17 have qualifying Week 24 source cells. 15 participants have 337 matched Week 24 target lineages, of which 332 enter the CDRH2 analysis (2,179 assessable lineage-position records, 241 observed state changes). The flow is about sampling and eligibility, not biological lineage loss or persistence.

#### 2.2.1. Longitudinal cell depth support and denominator variation

Our primary IGHV1-2*02 heavy chain CDRH2 analysis includes 2,179 eligible lineage-position observations from 332 lineages and 15 participants, with 241 observed acquisitions. Group 1 has 919 eligible observations, and Group 2 has 1,260. The median number of assessable cells per eligible lineage-position at each visit is 1 in both groups. If we require at least 2 assessable cells at each visit, we have 39/919 (4.24%) Group 1 observations and 100/1,260 (7.94%) Group 2 observations. If we require at least 3 cells per visit, we have 14 (1.52%) and 26 (2.06%) respectively. This descriptive depth assessment does not apply the additional modal frequency requirement as in the separate strict support sensitivity analysis. The original 7-position profiles are complete for all 15 participants. For the 2-cell threshold, complete profiles are maintained for 2/7 Group 1 participants and 4/8 Group 2 participants. For the 3-cell threshold, only 2 participants per group maintain complete profiles. These restrictions substantially alter the population evaluated. Thus, a meaningful like-for-like participant-level robustness comparison is not possible. Original eligible lineage denominators vary across participant-position combinations: 4–71 (median 9) in Group 1, and 5–56 (median 19.5) in Group 2. Despite these differences in denominator size, we equally weight each participant in our primary comparison. This sparse and uneven longitudinal cell support limits confidence that observed modal state differences are newly acquired biological substitutions.

#### 2.2.2. Sampling depth sensitivity

We explore cell-level support for modal state transitions at the 7 originally selected CDRH2 positions (51, 52, 53, 54, 55, 57, 58). For strict sampling support, we require at least 2 assessable cells at each visit and modal residue fraction *>* 0.5 at both visits. We have 3.25–4.50% eligible Group 1 observations and 6.43–8.89% eligible Group 2 observations with these criteria across the 7 positions. At least 1 strictly supported eligible lineage at each position is present for 2/7 Group 1 participants and 4/8 Group 2 participants. 9/15 participants have strictly supported observations at one or more positions. At position 55, we have 5/45 original Group 2 acquisition records with strict support (3 participants). No Group 1 acquisition is observed (either definition). This substantial reduction in observations, small number of lineages, and uneven participant coverage limit the feasibility and interpretability of a depth-restricted regional comparison. We do not treat the restricted observations as independent validation of the original regional profile result. We do not perform an additional confirmatory permutation analysis. This indicates that most observed modal state transitions have limited multi-cell longitudinal support. Thus, our primary analysis describes differences between sampled lineage states, not demonstrating mutation fixation, exact timing of mutation acquisition, or direct antigen-driven selection.

### 2.3. Paired probe sensitivity of observed lineage recovery

To evaluate antigen-dependent ascertainment, we examine the 7 original Group 1 CDRH2 participants with both eOD-GT8- and Cg28v2-sorted VRC01-class IGHG cells at Week 24. We keep frozen Week 8 lineage definitions and original unique assignment algorithm. We exactly reproduce all 2,202 Cg28v2 cell assignments in the corresponding frozen reference with the modified implementation. We have 181 recovered lineages (eOD-GT8-sorted) and 143 recovered lineages (Cg28v2-sorted) among 1,846 IGHV1-2*02 baseline lineages. We detect 60 lineages in both probe-sorted samples. We detect 121 lineages only in eOD-GT8 and 83 lineages only in Cg28v2. This establishes partial discordance in sampled lineage recovery. But we cannot attribute this solely to antigen binding specificity given sparse detection and different cell depths. The participant-equal mean difference in recovered baseline lineage fractions (Cg28v2 minus eOD-GT8) is *−*2.72 percentage points. The mean difference after 1,000 repeated equal cell depth samples within participants is *−*0.76 percentage points. The decrease in contrast after cell depth equalization indicates unequal sampled cell counts contribute to the numerical difference in recovery fractions. This resampling does not test equal depth overlap of *lineage identities*. Low overlap may also result from sparse sampling. The range of resampling is about variation within the observed cell pools, not a confidence interval for the participant population. Detailed paired probe findings are reported in the Supplementary Methods and Results.

We also explore the feasibility of estimating probe-specific CDRH2 positional state profiles. We recover 60 baseline lineages with both Week 24 sorting probes. These provide 214 uniquely assigned cells (105 eOD-GT8, 109 Cg28v2). A feasibility criterion (fixed prior to inspection of position eligibility) is at least 5/7 participants have at least 5 shared eligible lineages at every one of the 7 CDRH2 positions. All 420 possible shared lineage/position combinations have potential amino acid coverage under both probes. Shared lineages are unevenly distributed across participants (0, 3, 3, 19, 2, 2, 31). Only 2/7 participants can satisfy this liberal coverage-only support threshold. Given 5 qualifying participants are required, we do not perform the stricter eligibility assessment and positional probe sensitivity statistic.

### 2.4. Secondary prediction of observed Week 24 lineage recovery

We evaluate observed recovery at Week 24 for 2,739 Week 8 lineages from 17 participants in a separate Group 2 prospective benchmark. We detect 243 lineages (8.87%) at follow-up. Four participants have no observed recovered lineages. We must not conflate this binary sequence detection outcome with the 15 participant CDRH2 modal residue acquisition endpoint. We evaluate 5 fixed benchmark models on the same lineages and participant-held-out folds (Figure 3). The non-sequence biological baseline achieves participant-balanced AP of 0.107994 and participant-balanced AUROC of 0.615354 (Table 1). The participant-balanced AP null reference is 0.061165. The pooled outcome prevalence is 0.088719. We do not interchange these distinct reference quantities. Sequence-only IgBERT and AbLang-2 models have participant-balanced AP of 0.093925 and 0.081484 respectively. Baseline + IgBERT and baseline + AbLang-2 have participant-balanced AP of 0.096700 and 0.093614 respectively.

**Figure 3:**
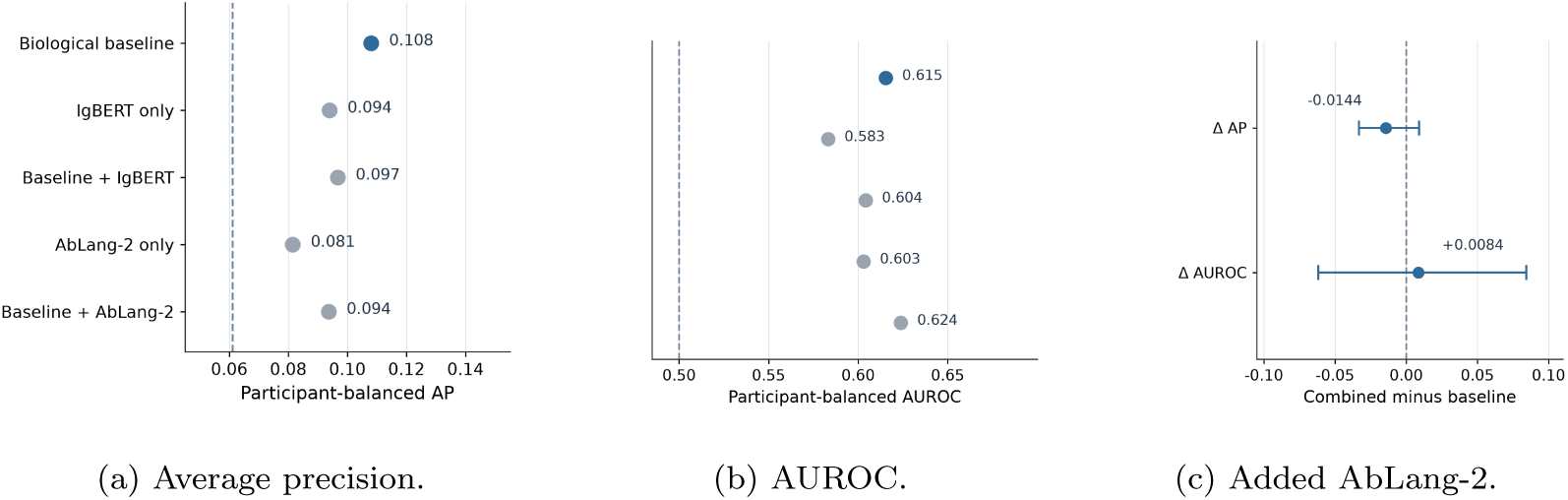
Observed Week 24 lineage detection prediction. Group 2. 2,739 Week 8 lineages. 17 participants. (A) Participant-balanced average precision (AP). Dashed reference is balanced outcome prevalence (0.061165), not pooled prevalence (0.088719). (B) Participant-balanced AUROC. (C) Baseline + AbLang-2 minus biological baseline. ΔAP = *−*0.014380 (95% participant-cluster bootstrap interval: *−*0.033168 to 0.008914). ΔAUROC = 0.008419 (interval: *−*0.061794 to 0.084374). Intervals describe uncertainty in frozen held-out predictions, not externally validated operational performance.

**Table 1.** Frozen participant-balanced prediction performance for observed Week 24 lineage recovery.

| Model | Participant-balanced<br>AP | Participant-balanced<br>AUROC |
|---|---|---|
| Biological baseline | 0.107994 | 0.615354 |
| IgBERT sequence only | 0.093925 | 0.583141 |
| Baseline + IgBERT | 0.096700 | 0.604094 |
| AbLang-2 sequence only | 0.081484 | 0.602925 |
| Baseline + AbLang-2 | 0.093614 | 0.623773 |

For baseline + AbLang-2 versus baseline, the reported difference in participant-balanced AP is *−*0.014380 (95% paired participant-cluster bootstrap percentile interval: *−*0.033168 to 0.008914). Difference in participant-balanced AUROC is 0.008419 (95% interval: *−*0.061794 to 0.084374). Both intervals include zero. Under this specific cohort, outcome, feature construction, and validation protocol, the tested antibody sequence embeddings do not show incremental improvement in participant-balanced AP.

## 3. Discussion

Incremental contribution and interpretation: The new contribution beyond Willis et al. [1] is an exploratory participant-balanced comparison of within-CDRH2 positional acquisition patterns between the two G002 vaccination groups. Participant-label permutation testing, leave-one-position-out sensitivity analyses, and descriptive sampling-support and probe-ascertainment assessments were used for the regional comparison. These results do not indicate timing or fixation of mutations; nor do they indicate vaccine-induced causality at any given residue, or antigen affinity/neutralization improvement. Interpretation is limited by sparse longitudinal sampling, probe-related unresolved ascertainment, position selection from data, and residue assessability. Thus, the result is a testable descriptive pattern that adds to, rather than replaces, the original trial’s functional findings. This secondary reanalysis reveals a difference between regimens in the shape of observed CDRH2 lineage-modal-state transition profiles, rather than directly testing their mean burden.

The participant-balanced seven-position comparison of 15 participants showed an exact permutation p-value of 0.001709 (six-region adjusted q = 0.010256). For Group 1, there is a higher descriptive acquisition fraction at V-relative position 51, while for Group 2, there are higher fractions at 54 and 55. These individual positions were not individually confirmed by inferential tests. The absence of observed acquisition at position 55 for Group 1 is a descriptive observation, not a structural biological zero or individually tested position-specific effect. To determine whether this is due to biological differences, differential ascertainment, or both, comparable sorting and longitudinal sampling would be required.

The current analysis focuses on a different longitudinal estimand than the original G002 report. Willis et al. already demonstrated regimen-associated differences in HCDR2 key-residues, showed position-specific residue patterns, and discussed development in selected large lineages [1]. Our contribution is a participant-level comparison of matched Week-8-to-Week-24 lineage-modalstate transition profiles, along with reconstruction and ascertainment sensitivity assessments. This distinction in analytical estimands is supported by comparison with their Figure 5E–F, Figure S39, and Table S61; this is not to say that all supplementary analyses have been exhaustively ruled out as precedents. For seven Group-1 participants with paired Week-24 probesorted samples, 181 prospective baseline lineages were recovered with eOD-GT8 sorting, 143 with Cg28v2 sorting, and 60 with both. When sampling depth was equalized, the contrast in participant-equal recovery-fraction was reduced from 2.72 percentage points to 0.76 percentage points. Thus, the observed contrast in recovery is sensitive to equalization of sampling depth, while the sets of recovered lineages are only partially overlapping. In a separate Group-2 prediction cohort, IgBERT and AbLang-2 did not improve participant-balanced average precision over a four-feature biological baseline when representations were added with leave-one-participant-out validation. This bounded negative result is for observed Week-24 lineage recovery and does not explain the mechanism or provide independent validation of the CDRH2 association. These tested representations must be interpreted in light of published antibody-language-model development and evaluation [7, 8], and prior work on vaccination-associated B-cell receptor repertoire prediction using a different experimental endpoint [9].

### 3.1. Limitations

Lineage-state changes are changes in the most frequently sampled aminoacid residue, not directly timed/fixed mutations. Changes in sampled subclones may lead to different modal states. V-relative positions were not functionally mapped to experimentally characterized antibodies. Thus, positional contrasts do not demonstrate improved binding/neutralization, or favorable mutation effects. Longitudinal sampling is sparse: for 2,179 eligible lineage-position observations, the median assessable cell count is one per visit. For at least two cells at both visits, 4.24% of Group-1 observations and 7.94% of Group-2 observations are retained. For three, 1.52% and 2.06% are retained, respectively. This reduces participant support too much for an equivalent confirmatory regional comparison. Thus, the exact permutation result is for the original definition of assessability, not for deeply sampled subclonal histories.

Both regimen groups are defined by eOD-GT8-sorted Week-8 cells and Cg28v2-sorted Week-24 cells. Thus, the endpoint is conditional on antigen-sorted lineage recovery and follow-up assessability. For Group 1, equal-depth paired-probe resampling addressed sampled cell numbers, but does not provide equivalent recovery of lineage identities. A separate attempt to assess probe-specific positional states was stopped at the feasibility gate fixed prior to position-level eligibility inspection. Only two out of seven Group-1 participants could meet the liberal requirement of five shared lineages with potential amino-acid coverage at all seven positions, while five qualifying participants were required. The stricter Week-8 and Week-24 state-eligibility criteria were not evaluated. No positional probe-effect statistic was estimated. For Group 2, there are no corresponding paired Week-24 sorting observations. Thus, the role of sorting-probe ascertainment in the observed positional difference between regimens remains unresolved. Finally, while the acquisition-support screening was invariant to participant-label relabeling, the overall analysis strategy was not prospectively registered. The exact test is conditional on the selected assessable participants and participant-label exchangeability; six-region multiplicity correction does not account for all exploratory analytical choices. Participant-omission and position-omission checks are internal robustness checks, not external cohort replication.

The data support a participant-resolved difference in the observed positional distribution of sampled CDRH2 lineage states. To establish the underlying biological mechanism, more deeply sampled paired repertoires, comparable antigen-sorting strategies across groups, and linked functional antibody measurements would be required. Biological context on the importance of VRC01-class CDRH2 residues is provided by published experimental work. Chen et al. (2021) [11] studied antibody substitutions at Kabat positions 54 and 57 in a mouse model. For the verified crosswalk to IGHV1-2*02 (Supplementary Table S3), these are V-relative positions 55 and 58, respectively. Between studies, reference sequence, antibody backgrounds, and substitutions must be distinguished. Coordinate correspondence alone does not indicate equivalent residue effects. In the current G002 analysis, there is no matched functional test for observed lineage-modal substitutions. Thus, neutralization/binding effects cannot be attributed to these substitutions.

An additional exploratory hierarchical sensitivity analysis is reported in Supplementary Section 9; its shrinkage-based estimates do not independently confirm or refute the participant-balanced permutation result.

## 4. Methods

### 4.1. Baseline lineage construction

Baseline cells were selected from the provided G002 table for Groups 1 and 2, Week 8, probe_set eODGT8, VRC01-class status, and constant-region call IGHG. Clustering was performed within each participant. Existing cluster labels and centroid labels were removed. A SADIE LinkedAirrTable keyed by cellid was then constructed. The SADIE Cluster configuration used average linkage and grouping variables: heavy-chain V call, light-chain V call, heavychain junction amino-acid length, constant-region call, and VRC01-class status. Heavy- and light-chain CDR3 amino-acid sequences were provided as lookup fields. Somatic padding was disabled (pad_somatic=False). Clustering was invoked with model.cluster(3). Prospective lineage identifiers were created by concatenating participant identifier and cluster identifier. Later-visit cells were not included in this baseline clustering. Baseline clustering and follow-up assignment had different rules. For follow-up assignment, compatible heavy/light J calls and light-chain junction length were also required. The mean summed CDR3 edit-distance threshold was four. Reconstructed cluster membership was compared to the provided author annotation on shared cell identifiers. The adjusted Rand index and exact cell-level membership agreement were used.

### 4.2. Prospective follow-up matching

Baseline lineage identifiers were read from the saved Week-8 table. If missing, baseline lineage identifiers were created by concatenating participant identifier and cluster identifier. Separate documentation was provided for baseline clustering. Prospective matching was performed on follow-up records separately for Weeks 16 and 24. Eligible records were restricted to Groups 1 and 2, probe_set Cg28v2, VRC01-class status, and top constant-region call IGHG. The primary lineage-state transition analysis was for Week-8 states versus uniquely assigned Week-24 cells. Week-16 records were not included in this endpoint. Candidate baseline lineages were restricted to the same participant, heavy- and light-chain V calls, heavy- and light-chain junction lengths, and VRC01-class status. Later heavy- and light-chain J calls must also equal the corresponding baseline lineage modal J calls. Baseline V calls, junction lengths, and class status were represented by the first recorded baseline cell per lineage. For heavy- and light-chain J calls, the nonmissing modes were summarized. If tied, the first value returned from the mode calculation was used. For each candidate lineage, the heavy-chain CDR3 amino-acid Levenshtein distance and light-chain CDR3 amino-acid Levenshtein distance were summed for each baseline cell. These summed distances were then averaged across baseline cells. A lineage was qualified if the mean distance was at most four. A later cell was assigned only if exactly one lineage qualified. If no lineage qualified, the later cell was classified as unmatched. If multiple lineages qualified, the classification was ambiguous. The cell was not assigned to the closest lineage. At each participant-visit, assessability was defined as having at least one later cell meeting the above later-record filters. For assessable visits, a lineage was detected if at least one later cell was uniquely assigned to the lineage. For visits with no eligible later cells, the detection status was missing, rather than negative. Thus, detection is dependent on sampling and the specified matching rule. It does not indicate biological persistence or disappearance.

### 4.3. Within-lineage sequence states and mutation transitions

#### 4.3.1. Follow-up records

Follow-up residue states were constructed from Week-24 cells uniquely assigned to baseline lineages. Week-8 residue states were only evaluated for lineages represented among these assigned Week-24 cells. This used the existing lineage assignments created during the Week-8 baseline clustering and prospective follow-up matching procedures, as described above.

#### 4.3.2. Assessable residues

For each cell and chain, a local V-region amino-acid mutation fraction was calculated. This was the number of observed-versus-germline residue mismatches divided by the number of assessable residues in the paired V-region alignment. Gap and ambiguous (X) residues were excluded from this local calculation. The absolute validation difference was the absolute difference between this local fraction and the corresponding source-annotated v_mutation_aa_heavy or v_mutation_aa_light fraction. Cell-chain records were only retained if there was no gap in the observed or germline V-region alignment, and the absolute validation difference was no greater than 0.001. Positions were numbered from one in the paired V-region amino-acid alignment strings. Positions with X in either the observed or germline sequence were excluded.

#### 4.3.3. Within-lineage states

Residue counts were calculated for each combination of lineage, chain, V-gene call, position, and germline residue at each visit. One residue observation was contributed by each assessable cell. The observed lineage state was the most frequent residue. If two or more residues tied for the highest count, the state was unresolved. No additional minimum frequency or strict-majority threshold was used.

#### 4.3.4. Transition classification

Week-8 and Week-24 states were joined on lineage, chain, V-gene call, position, and germline residue. This was a one-to-one merge requirement. Only keys represented at both visits were included in the transition table. If either visit had a missing state, the transition was classified as unresolved. If the Week-8 state was germline modal, and the Week-24 state was non-germline modal, the transition was classified as acquired_from_germline. Other classifications included stable germline states, unchanged preexisting mutations, reversions to germline, and changes between different non-germline states.

#### 4.3.5. Biological interpretation

The acquisition endpoint is a germline-to-non-germline change in the observed within-lineage modal residue between visits. This does not indicate that the non-germline residue was absent from all Week-8 cells, arose after Week 8, or became fixed in the lineage. Changes in the frequency of sampled subclones may also change the modal state. Interpretation is still conditional on sampling, lineage assignment, and residue assessability.

### 4.4. Regional lineage-state transition profile methods

#### 4.4.1. Eligible observations

The regional analysis used the saved Week-8-to-Week-24 mutation-transition table. Records were restricted to heavy-chain IGHV1-2*02 observations, with a transition classification other than unresolved. Week-24 amino-acid state must not be missing. Week-8 state must equal the germline state. Acquisition was defined as a Week-24 state different from germline. For each participant and position, the acquisition fraction was calculated as the sum of acquisition indicators divided by the number of distinct eligible lineages. To interpret this as a lineage fraction, there must be one eligible record per lineage and position. These fractions describe changes in recorded sequence states. They do not indicate the timing of a mutation or describe undetected lineages.

#### 4.4.2. Original position selection

Position inclusion was first evaluated using pooled acquisition support within (position, germline amino-acid residue) combinations. Positions with at least five acquired lineage observations spanning at least three participants were retained, with support pooled across groups. Position columns with nonmissing fractions for all analyzed participants were included in the regional comparison. Regions were assigned using the saved coordinate-region mapping. The original CDRH2 comparison was for seven V-relative positions: 51, 52, 53, 54, 55, 57, and 58. The positions are based on the ungapped IGHV1-2*02 germline V-region sequence using 1-based coordinates. The germline residue identities are cross-referenced with the reference accession X62106. Independently, Kabat numbering was evaluated with ANARCII for 23 gap-free G002 heavy-chain sequences across 14 participants. 161 evaluated residue assignments are in agreement. Similarly, the independent IMGT numbering of the X62106 germline V-region reference using ANARCII v2.0.8 are in agreement for all 7 positions. The verified coordinate correspondence is shown in Supplementary Table S3. The purpose of these evaluations is to ensure numbering correspondence for the assessed reference and sequences, not functional cohort-wide validation. At the original acquisition-support screening step, the V-relative position 56 was excluded. Only one lineage from one participant showed observed germline-to-nongermline transition at this position (below the 5 acquisition and 3 participant thresholds) among the eligible IGHV1-2*02 heavy-chain records. The subsequent coordinate-defined exploratory sensitivity analysis for positions 51-58 (same cohort) is not an independent validation.

#### 4.4.3. Analysis-selection chronology and inferential scope

The original acquisition-support filtering pooled observations across the vaccination groups. Participant-group labels were not used. This screening rule is dependent on the pooled observed lineage-state data, not on the group assignments. Thus, its selection outcome is invariant to the permutations of the participant-group labels, given the fixed observed data. Therefore, the same positions would be selected if the identical screening rule was reapplied for each participant-label permutation. This is relevant to the distinction between label-invariant filtering and group-informed selection discussed in statistical literature [5,6]. However, it does not imply independence of the screening criterion and test statistic for all null models or remove the selection-related concerns. An exact participant-label permutation test was used to assess the original 7-position comparison for CDRH2. For 15 participants (7 Group 1 and 8 Group 2), all 6,435 label allocations were enumerated. The observed centered-profile statistic was 0.027066. 11/6,435 allocations showed a statistic equal to or larger than the observed value (p = 0.001709). The reported permutation p-value is for the implemented comparison conditional on the selected participant profiles. Its interpretation is dependent on the participant-label exchangeability under the null within the analyzed cohort. It is not implied that there are no differences in lineage detection, follow-up assessability, or antigen-probe ascertainment. While the pooled position-selection rule was label-invariant, the reviewed planning records did not show prospective specification of the complete analysis strategy, including the endpoint, regional focus, and the screening thresholds. Thus, the CDRH2 comparison is presented as exploratory rather than confirmatory. The Benjamini-Hochberg adjustment was applied across the original 6 regional comparisons (CDRH2 adjusted q = 0.010256). This does not account for the broader history of exploratory analysis choices. The subsequent analyses (same participants) are not independent validation.

#### 4.4.4. Centered-profile statistic

For each participant, the mean of the acquisition fractions across the included positions of a region was subtracted from the position-specific fractions. The resulting centered fractions were then averaged across participants within each group. The mean squared difference between the 2 group-average centered profiles across positions was then calculated to give a measure of the shape of the profile after removing the regional mean for each participant. This is not a direct test of the overall mutation burden.

#### 4.4.5. Permutation inference and multiplicity

The participant group labels were permuted. The observed group sizes were preserved. All 6,435 allocations were enumerated for 15 participants (7 Group 1). The p-value was the proportion of allocations with the statistic greater than or equal to the observed statistic. The squared statistic detects differences in profile in either direction. Benjamini-Hochberg adjustment was applied across the original 6 regional comparisons. It was not applied for the subsequent omission, 8 positions, or structural analyses. The inference assumes exchangeability of the participant profiles under the null within the analyzed population. Follow-up assessability is a limitation.

#### 4.4.6. Descriptive cell-depth and denominator assessment

We evaluated the original 7 CDRH2 positions in the original 15-participant IGHV1-2*02 analysis cohort using the frozen Week 8 to Week 24 transition records. For each eligible lineage-position record, the number of assessable cells contributing to the Week 8 and Week 24 residue states were extracted from the saved transition table. Separate minimum assessable-cell thresholds of 1, 2, and 3 were applied for both visits. The original positions and participants were used as the reference population. The positions were not reselected. For each threshold, we summarized the retained lineage-position observations, acquisition counts, denominators for eligible lineages, and the number of participants retaining all 7 position-specific fractions. For the missing (rather than 0) fractions (where a participant did not have an eligible lineage at a position), this was a descriptive measurement-support analysis. It is not an independent validation or a new permutation test. Unlike the separate strict-support analysis, the thresholds did not require a modal residue fraction > 0.5.

#### 4.4.7. Sampling-support sensitivity

A post hoc sensitivity analysis was performed to assess sampling support for the mutation transitions at the 7 original CDRH2 positions. For each eligible lineage-position record, the strict support was at least 2 assessable cells for Week 8 and Week 24, and modal amino-acid fractions > 0.5 for both visits. For each participant and position, the number of eligible lineages, acquired mutations, strictly supported eligible lineages, and strictly supported acquisitions was counted. The complete 7-position support was defined as at least 1 strictly supported eligible lineage for each of the 7 positions for a participant. The positions could be supported by different lineages for the same participant. The results were summarized at the participant and group levels. No new confirmatory regional permutation test was performed on the restricted subset. The strict criterion was used to assess measurement support and feasibility. It is not for establishing mutation fixation or biological selection. Strict restriction may also introduce selection bias due to varying cell sampling depth and lineage eligibility across participants and lineage characteristics.

### 4.5. Within-visit probe-ascertainment sensitivity

We evaluated the 7 Group 1 participants included in the primary CDRH2 analysis using their frozen Week 8 eOD-GT8 lineage definitions and Week 24 eOD-GT8- or Cg28v2-sorted VRC01-class IGHG cells. Both the probesorted populations were analyzed using the original compatibility criteria, mean summed heavy/light CDR3 Levenshtein distance threshold of 4, and unique-only assignment rule. The existing Cg28v2 assignment results were used as a computational replay reference. Recovery was defined as at least 1 uniquely assigned Week 24 cell from a prospective IGHV1-2*02 Week 8 lineage under the specified probe. The same baseline-lineage denominators were used for the participant-specific recovery fractions across both probes. To describe the sensitivity to the sampling depth, 1,000 repetitions were performed to sample the same number of cells (without replacement) from the 2 probe-sorted populations for each participant. The smaller observed cell count was used as the target depth. The differences were averaged with equal weight for the participants. The distribution of repeated equal-depth estimates was treated as a conditional Monte Carlo sensitivity distribution, not as the inferential uncertainty across participants. No new hypothesis tests were performed.

#### 4.5.1. Paired-probe positional-state feasibility

The separate positional-state feasibility assessment used 60 baseline lineages that were recovered by both probes for the 7 original Group 1 participants. Prior to examining the eligibility at the position level, a decision rule was applied. It required at least 5 participants, each with at least 5 shared eligible lineages for each of the selected CDRH2 positions (51, 52, 53, 54, 55, 57, 58). Full eligibility was defined as having a resolved germline Week 8 modal state and separately assessable Week 24 modal states under both probes (as per the original sequence-validation and tie rules). A deliberately liberal necessary-condition assessment was used. It only required the non-X amino-acid coverage under both probes. All 60 shared lineages were retained regardless of the Week 8 state. If this upper-bound support criterion failed, the planned positional comparison was terminated without estimating the statistic or changing the eligibility threshold. This feasibility rule was fixed for the exploratory sensitivity investigation. It was not prospectively registered with the original vaccination trial.

### 4.6. Secondary prediction cohort, features, and validation

The separate Group 2 prospective prediction cohort had 2,739 Week 8 lineages (17 participants). 243 were detected at Week 24. 5 models were used with the same outcome labels, participants, and leave-one-participant-out folds. The 4-feature non-sequence baseline was the Week 8 log2 lineage cell count, heavy-chain somatic hypermutation, light-chain somatic hypermutation, and Cottrell mean. IgBERT used 1,024-dimensional frozen sequence-medoid representations. AbLang-2 used 480-dimensional frozen paired heavy- and light-chain mean-pooled representations. L2-regularized logistic regression (C = 1) was used. Scalers (training only) and equal-participant sample weights were used. No PCA was used. Participant-balanced and pooled metrics were used for the benchmark, but without conflating their AP prevalence references. Previously saved participant-cluster bootstrap summaries (2,000 resamples of paired out-of-fold predictions) were used for the reported differences and 95% intervals. They were not recomputed during manuscript preparation. Full technical details and complementary performance metrics are retained in the supplementary Methods and Results.

## Supporting information

Supplementary Methods and Results

## 5. Data and Code Availability

### 5.1. Data availability

This independent secondary computational study analyzed previously published IAVI G002 B-cell receptor repertoire data reported by Willis et al. [1]. The original source data and associated analysis materials can be accessed from the original investigators at:

https://github.com/SchiefLab/G002-and-G003

The archived release of the original study is available at:

https://doi.org/10.5281/zenodo.15425848

The original final_df.feather source file used for this reanalysis was previously verified to be byte-identical to the author-released corresponding source file. Fourteen selected derived aggregate result tables (regional comparisons, sensitivity analyses, secondary prediction summaries, and 7-position Figure 1A group-mean table) are publicly available at: https://github.com/KaziTanzina/HIV-Vaccine-BCR/tree/main/data/ aggregate_release_candidate

The exact documented public release revision is commit c6ee284 of that repository. The original third-party sequence-level records and individual participant-level plotting observations are not included in the released aggregates. Thus, the aggregate tables alone are not sufficient to reproduce all the participant points in Figure 1A. They are not a complete end-to-end reproduction dataset.

### 5.2. Code availability

The G002 and G003 analysis software of the original investigators is available at the SchiefLab repository (cited in Data Availability).

The computational code for the present independent secondary reanalysis was developed by Kazi Tanzina Begum. It is publicly available at: https://github.com/KaziTanzina/HIV-Vaccine-BCR

A fixed reference for the publicly released materials is: https://github.com/KaziTanzina/HIV-Vaccine-BCR/commit/c6ee284 The earlier collection of 22 selected analysis scripts and 4 publication figures are in the repository. A separate package of 11 checksum-verified primary-analysis and Figure 1A reconstruction scripts (which may overlap with the earlier script collection) is also available. The 14 derived aggregate tables are also included. The study-specific Python code is distributed under the BSD 2-Clause software license. The separate provenance and applicable reuse conditions apply for the original third-party data and software. The participant-level Figure 1A reconstruction and 65-position centering calculation were verified against the frozen research outputs. However, there has been no independent validation of the published materials as a complete end-to-end pipeline (starting from a fresh download of the original G002 data). Some required frozen intermediate files (including the participant-level plotting inputs) are not redistributed. The selected primary reproducibility package does not include a standalone rerun of all the historical antibody-language-model embedding-generation steps.

## 6. Declarations

### 6.1. Ethics approval and consent to participate

This independent computational secondary study reanalyzed previously published IAVI G002 B-cell receptor repertoire data. No new participants were recruited. No additional biological specimens were collected. No new experimental interventions were performed. Willis et al. [1] reported written informed consent and original trial institutional review board approvals at George Washington University (NCR213652), University of Texas Health Science Center at San Antonio (21-339H), Fred Hutchinson Cancer Center (10723), and Emory University (STUDY00002864). These approvals are for the original clinical investigation. They are not represented as separate approval for this independent secondary analysis. There is no separate institutional review board determination for the present study.

### 6.2. Consent for publication

Not applicable. This secondary computational analysis does not present identifiable individual case material or participant images. Separate consent for publication is not required.

### 6.5. Author contributions

Kazi Tanzina Begum led the study design. She developed the analytical framework and performed the computational analyses. She developed the software and prepared the manuscript. Kazi Abdul Gofur Rafea contributed to the study design and methods. Both authors reviewed and approved the final manuscript for submission.

### 6.6. Declaration of generative AI use

We used OpenAI to assist with code debugging. We reviewed and revised the content. We also checked the computational results against the underlying data and the saved analysis outputs. We take full responsibility for the scientific content, analyses, interpretations, and conclusions.

### 6.7. Acknowledgments

No additional acknowledgments.

