## Supplementary Methods and Results for "Exploratory positional differences in observed CDRH2 lineage-state transitions following HIV-1 germline-targeting vaccination"

### Supplementary Methods and Results — Repertoire sampling, prediction benchmarks, and CDRH2 residue numbering

---

---

#### 1. Scope

This supplement covers the six aspects of the G002 secondary analysis: (i) the depth of sampling required for the primary CDRH2 analysis; (ii) the sensitivity of detecting lineages with the same-visit paired-probe; (iii) the feasibility of a probe-specific CDRH2 positional-state sensitivity; (iv) an exploratory hierarchical modeling sensitivity; (v) a separate Group-2 antibody-language-model prediction benchmark; and (vi) the independently verified IGHV1-2*02 CDRH2 residue-numbering crosswalk. Note that the prediction endpoint for this analysis is observed Week-24 lineage detection, not the acquisition of mutations or antibody function.

#### 2. Group-2 prediction benchmark — Supplementary Methods

##### *2.1. Prediction cohort and outcome*

Frozen analysis included 2,739 prospective lineages from 17 participants at Week 8. Of these, 243 were detected at Week 24. Prediction outcome was frozen sequence-matching and repertoire sampling procedure-based binary detection. Nondetection does not necessarily imply biological loss of a B-cell lineage.

##### *2.2. Biological baseline*

The original frozen biological comparator was composed of four Week-8 features:

1. Log2 Week-8 lineage cell abundance.
2. Heavy-chain somatic hypermutation percentage.
3. Light-chain somatic hypermutation percentage.
4. Mean Cottrell score.

##### *2.3. Pretrained antibody-language-model representations*

IgBERT and AbLang-2 were tested on previously saved pretrained representations of sequences. The IgBERT representation used for frozen analysis was of size 1,024. AbLang-2, with the pretrained paired heavy/light-chain encoder (package version 0.1.1), was used with existing observed Week-8 paired-chain medoid sequences. New representatives were not chosen for the representation. The amino acid positions on the heavy and light chains, excluding the separator and padding positions, were averaged to obtain a residue-level representation of 480 features per lineage.

The pretrained encoders were not fine-tuned on the G002 outcome labels.

##### *2.4. Prediction and validation*

Models were defined as the biological baseline, each of the sequence-only representations, and the combination of each representation with the four baseline predictors.

The same 2,739 lineages, outcomes and participant-held-out folds were used for all comparisons. Leave-one-participant-out was used for cross-validation, i.e. the held-out participant was not used for fitting the classifier. The classifier used in the frozen analysis was an L2-regularized logistic regression model with C = 1.0, liblinear solver, balanced class weights, and training-participant-equalizing sample weights. Standardization was fit on each fold and then applied to the held-out fold. No principal-component reduction or additional feature selection was applied to the pretrained embeddings.

##### *2.5. Evaluation metrics and prevalence references*

Out-of-fold discrimination was measured by AUROC and average precision (AP) (pooled and participant-balanced).

Participant-balanced discrimination was assessed by weighting each lineage by the reciprocal of the total number of eligible lineages for that participant, i.e. each participant was given equal weight.

The pooled outcome prevalence is **0.088719**. The participant-balanced prevalence is **0.061165**, i.e. the average of the 17 participant-specific detection fractions. These prevalences correspond to the AP of an uninformative constant-score predictor. They are given as descriptive reference values, not as significance tests by permutation.

##### 2.6. Uncertainty

We used 2,000 paired participant-cluster bootstrap resamples to evaluate the frozen comparison of baseline plus AbLang-2 versus the biological baseline. The predictions and the observed labels were kept paired. The percentile 95% intervals indicate the uncertainty in the saved out-of-fold performance comparison given the fitted models and the completed model-development procedure.

#### 3. Group-2 prediction benchmark — Supplementary Results

##### 3.1. Model comparison

| Model | Pooled AUROC | Pooled AP | Balanced AUROC | Balanced AP |
|---|---|---|---|---|
| Biological baseline | 0.618722 | 0.144932 | 0.615354 | 0.107994 |
| IgBERT only | 0.568772 | 0.114091 | 0.583141 | 0.093925 |
| Baseline + IgBERT | 0.595334 | 0.124632 | 0.604094 | 0.096700 |
| AbLang-2 only | 0.577477 | 0.111838 | 0.602925 | 0.081484 |
| Baseline + AbLang-2 | 0.594695 | 0.130622 | 0.623773 | 0.093614 |

*Balanced* metrics give each participant equal weight.

The corresponding AP prevalence references were **0.0887** for the pooled evaluation and **0.0612** for the participant-balanced evaluation.

The biological baseline had a participant-balanced AP of **0.1080** and the corresponding prevalence reference of **0.0612**. The baseline had a participant-balanced AUROC of **0.6154**. This indicates modest discrimination but does not establish any operational or clinical prediction utility.

Neither of the language-model representations improved upon the biological baseline in terms of average precision when added to the frozen baseline. The baseline plus IgBERT had an AP of **0.0967** and the baseline plus AbLang-2 had an AP of **0.0936** compared to the **0.1080** of the frozen baseline. For baseline plus AbLang-2 minus the biological baseline, the participant-balanced AP difference is **-0.014380** (95% participant-cluster bootstrap interval **-0.033168 to** +**0.008914**). The corresponding participant-balanced AUROC difference is +**0.008419** (95% interval **-0.061794 to** +**0.084374**). The upper and lower bounds of both intervals include zero, therefore the analyses do not establish any performance gain or a statistically significant performance decrease when adding AbLang-2.

##### *3.2. Interpretation and limitations*

For the fixed evaluation protocol used, the antibody language model representations tested did not provide any incremental value in average precision above the four features of the biological baseline. The absolute level of discrimination of the biological baseline was relatively low. No model was found to be suitable for operational prediction of vaccine responses. There were only 17 independent participants in the analysis. There was substantial variation in the detected lineage prevalence between participants, and for several participants no lineages were detected at Week 24. Also the endpoint was conditional on the repertoire sampling, sorting, matching and sequence ascertainment. The results do not indicate that antibody sequences are uninformative biologically, that other language model architectures are not effective, or that undetected lineages have disappeared biologically. The tested frozen representations, the choice of embedding extraction and the linear classifier used for prediction represent only a small part of the possible modeling design space. The uncertainty of the performance on held-out predictions was assessed using saved predictions rather than re-running the model development process.

##### *3.3. Relation to published work*

The results of this study are in line with other studies on pretrained antibody and protein language models, finding that they can be very effective on some antibody tasks but provide less added value on other tasks. The published tasks for IgBERT and IgT5 were developed and benchmarked for the purpose of antibody engineering. The published task for AbLang-2 is the representation of antibody sequences and the suggestion of non-germline mutations. None of these tasks have been shown to be superior for the prediction of longitudinal HIV vaccine-lineage detection. Prior evaluations have shown the importance of model choice, embedding layer, validation strategy and simple baselines. Vaccine-associated BCR clonotype prediction has also been studied previously in a Tdap vaccination context, but the endpoint G002 is different. Thus, we do not claim that antibody-language-model-based prediction of vaccine-induced B-cell responses is novel.

#### 4. Selected literature


1. Olsen TH, Moal IH, Deane CM. Addressing the antibody germline bias and its effect on language models for improved antibody design.

*Bioinformatics*. 2024;40(11):btae618. https://doi.org/10.1093/bioinformatics/btae618
2. Kenlay H, Dreyer FA, Kovaltsuk A, Miketa D, Pires D, Deane CM. Large scale paired antibody language models. *PLoS Computational Biology*. 2024;20(12):e1012646. https://doi.org/10.1371/journal.pcbi.1012646
3. Deutschmann et al. Do domain-specific protein language models outperform general models on immunology-related tasks. *ImmunoInformatics*. 2024;14:100036. https://doi.org/10.1016/j.immuno.2024.100036
4. Wang M, Patsenker J, Li H, Kluger Y, Kleinstein SH. Language model-based B cell receptor sequence embeddings can effectively encode receptor specificity. *Nucleic Acids Research*. 2024;52:548–557. https://doi.org/10.1093/nar/gkad1128
5. Kanduri C, et al. Profiling the baseline performance and limits of machine learning models for adaptive immune receptor repertoire classification. *GigaScience*. 2022;11:giac046. https://doi.org/10.1093/gigascience/giac046
6. Richardson E, Willemsen L, Shinde P, Nielsen M, Peters B. Is the vaccination-induced B cell receptor repertoire predictable? *ImmunoInformatics*. 2025;19:100057. https://doi.org/10.1016/j.immuno.2025.100057

#### 5. Frozen computational provenance

All model comparisons were reconstructed from the following previously saved prediction, summary and uncertainty files.

- `results/bcr/g002_g003/248_phase4a_frozen_baseline_lopo_predictions.tsv`
- `results/bcr/g002_g003/261_phase4a_igbert_sequence_only_lopo_predictions.tsv`
- `results/bcr/g002_g003/264_phase4a_baseline_plus_igbert_lopo_predictions.tsv`
- `results/bcr/g002_g003/phaseG_ablang2/573_phaseG_ablang2_sequence_only_lopo_predictions.tsv`
- `results/bcr/g002_g003/phaseG_ablang2/574_phaseG_baseline_plus_ablang2_lopo_predictions.tsv`

- `results/bcr/g002_g003/250_phase4a_frozen_baseline_lopo_summary.tsv`
- `results/bcr/g002_g003/phaseG_ablang2/575_phaseG_ablang2_lopo_summary.tsv`
- `results/bcr/g002_g003/phaseG_ablang2/578_phaseG_participant_bootstrap_summary.tsv`

All comparisons used the previously frozen model predictions and summaries. No additional encoder selection or model tuning was performed during manuscript preparation.

#### 6. Longitudinal cell-depth support

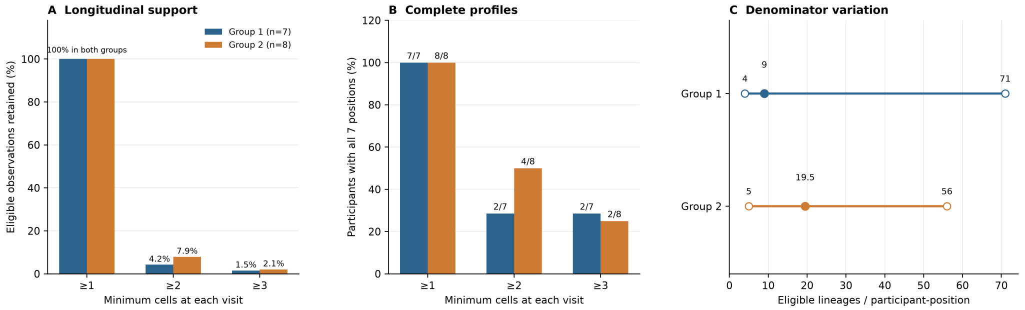


Figure S1: Longitudinal cell-depth support for the exploratory G002 CDRH2 analysis. (A) Percentage of eligible lineage-position observations retained with at least one, two or three assessable cells at both Week 8 and Week 24 (initially 919 Group-1 and 1,260 Group-2 observations). (B) Participant counts with complete seven-position coverage under those thresholds (Group 1, $n = 7$; Group 2, $n = 8$ at the original threshold). (C) Minimum, median and maximum original eligible-lineage denominators across participant-position records. The displayed cell-depth sensitivities are descriptive, not independent validation.

#### 7. Paired-probe Week-24 lineage-recovery sensitivity

This was a post hoc within-Group-1, within-visit analysis of seven primary CDRH2 participants with frozen Week-8 lineage definitions and original unique-assignment criteria. The Cg28v2 arm reproduced 2,202 frozen cell-level assignments exactly.

| Participant | Baseline lineages | eOD-GT8 recovered | Cg28v2 recovered | Both probes | eOD only | Cg only |
|---|---|---|---|---|---|---|
| G002-509 | 204 | 11 | 6 | 0 | 11 | 6 |
| G002-616 | 351 | 16 | 18 | 3 | 13 | 15 |
| G002-619 | 127 | 11 | 8 | 3 | 8 | 5 |
| G002-678 | 139 | 33 | 24 | 19 | 14 | 5 |
| G002-758 | 288 | 24 | 9 | 2 | 22 | 7 |
| G002-813 | 168 | 9 | 5 | 2 | 7 | 3 |
| G002-852 | 569 | 77 | 73 | 31 | 46 | 42 |

The recovery of 1,846 baseline target lineages was assessed under eOD-GT8 and Cg28v2 sorting probes at Week-24. For these probes, 181 and 143 lineages were recovered, respectively, and 60 of these were recovered under both probes. These results are based on observed recovery classifications and do not imply that any of these lineages had become biologically extinct.

The mean of the equal-participant Cg28v2-minus-eOD-GT8 recovery difference was -2.716 percentage points before depth equalization and -0.763 percentage points after 1,000 within-participant equal-depth samples. The 2.5th–97.5th percentiles of the Monte Carlo distribution are -1.464 to -0.094 percentage points. These values do not represent a population confidence interval.

Two populations of cells with different counts were compared in this analysis. The equal-depth analysis used to compare the two populations of cells with different counts assesses the differences in numbers of recovered-lineage between the two populations, not the identities of the recovered-lineages after equal-depth sampling. Thus, low observed overlap of lineages between the two populations does not imply that the two populations of cells have different antigen-binding specificities. This comparison is limited to Group 1, and cannot establish a causal booster effect or completely resolve the issue of ascertainment in the CDRH2 between-regimen comparison.

#### 8. Paired-probe positional-state feasibility — Supplementary Methods and Results

We first assessed whether the 60 shared lineages, recovered under both Week-24 sorting probes, were suitable for a probe-specific sensitivity analysis of the seven-position CDRH2 acquisition profile. First, we set a feasibility criterion, which required that for each of the 7 positions (51, 52, 53, 54, 55, 57, 58) in the seven-position CDRH2 acquisition profile, at least 5 of the 7 Group-1 participants had at least 5 shared eligible lineages. All 60 shared lineages had potential amino-acid coverage at all 7 positions (420 possible lineage-position combinations). The counts of shared lineages across the 7 participants were 0, 3, 3, 19, 2, 2, 31, respectively. Thus, only 2 participants had sufficient coverage at all 7 positions, which fell below the feasibility criterion of 5 participants. Hence, we did not evaluate full position-state eligibility, and we did not estimate any probe-dependent positional-profile statistic.

#### 9. Exploratory hierarchical sensitivity

We developed a post hoc Bayesian logistic model with random intercepts for participant and lineage as well as regularized interactions between group and position to reproduce the original cohort counts and total acquisition burden while producing smaller regional profile differences than the empirical analysis. The model-based posterior median regional statistic was 0.010839 whereas the empirical statistic was 0.027066. The model uses shrinkage priors for the group-by-position interactions, so the attenuation does not independently imply evidence against the original positional association. The model does not confirm or refute the permutation result based on participant-balanced permutations and does not allow for assessment of differential antigen-probe ascertainment.

#### 10. CDRH2 residue-numbering correspondence (Table S3)

V-relative positions are indexed from the ungapped IGHV1-2*02 germline V-region sequence (X62106). Germline identities were verified against this reference. Kabat assignments were checked on 23 gap-free G002 sequences from 14 participants (161 residue assignments). IMGT assignments were checked independently on the X62106 reference using ANARCII 2.0.8. The coordinate verification does not establish functional consequences of substitutions.

Table S3: Correspondence of IGHV1-2*02 CDRH2 residue numbering schemes.

| V-relative | Germline residue | Kabat | IMGT |
|---|---|---|---|
| 51 | I | H51 | 56 |
| 52 | N | H52 | 57 |
| 53 | P | H52A | 58 |
| 54 | N | H53 | 59 |
| 55 | S | H54 | 62 |
| 57 | G | H56 | 64 |
| 58 | T | H57 | 65 |
